# Plasmin cascade mediates thrombolytic events in SARS-CoV-2 infection via complement and platelet-activating systems

**DOI:** 10.1101/2020.05.28.120162

**Authors:** Kavitha Mukund, Kalai Mathee, Shankar Subramaniam

## Abstract

Recently emerged beta-coronavirus, SARS-CoV-2 has resulted in the current pandemic designated COVID-19. COVID-19 manifests as severe illness exhibiting systemic inflammatory response syndrome, acute respiratory distress syndrome (ARDS), thrombotic events, and shock, exacerbated further by co-morbidities and age^1–3^. Recent clinical reports suggested that the pulmonary failure seen in COVID-19 may not be solely driven by acute ARDS, but also microvascular thrombotic events, likely driven by complement activation^4,5^. However, it is not fully understood how the SARS-CoV-2 infection mechanisms mediate thrombotic events, and whether such mechanisms and responses are unique to SARS-CoV-2 infection, compared to other respiratory infections. We address these questions here, in the context of normal lung epithelia, *in vitro* and *in vivo*, using publicly available data. Our results indicate that plasmin is a crucial mediator which primes interactions between complement and platelet-activating systems in lung epithelia upon SARS-CoV-2 infection, with a potential for therapeutic intervention.

## MAIN

Recent studies^6,7^ have rapidly provided molecular insights into the pathogenicity of the SARS-CoV-2, mainly at the level of genomic, structural, and functional aspects of viral-host interactions. Further, studies have identified key pathophysiological and molecular events associated with infection pathogenesis and progression of COVID-19 including thrombocytopenia, lymphopenia, eosinophilia, and elevated lactate dehydrogenase and fibrinogen ^8,9^. The molecular findings have included a delayed interferon (IFN) response type I and III^10,11^ with a concomitant increase of proinflammatory cytokines (namely IL-17, IL-6 and/or IL-1B)^12^ leading to a “cytokine” storm coupled with the depletion of markers for platelets, natural killer cells, and dysregulation of CD^4+^ and B-cell lymphocyte populations ^13^. In contrast to other respiratory viral infections (e.g., ref^14–17^), SARS-CoV-2 can efficiently replicate in cells of various tissues that express angiotensin-converting enzyme 2 (*ACE2*) and host serine protease (*TMPRSS2*)^18,19^, thus contributing to the increased transmissibility and lower-lung pathogenicity in humans ^20^. This observation led us to explore mechanisms which are unique to SARS-CoV-2 infection of lung epithelial cells and cause thrombotic events.

In this study, we utilized publicly available RNA-sequencing data, GSE100457^10^, in Normal Human Bronchial Epithelial (NHBE) cell lines infected with SARS-CoV-2 (henceforth referred to as the CoV-2 dataset). In particular, we compared the CoV-2 dataset with lung epithelial cells infected with other respiratory infections, namely, respiratory syncytial virus (RSV), influenza (H1N1), and rhinovirus (RV16) (see Methods)), using a stringent study inclusion criterion. Differentially expressed genes (DEGs) were called for each infection (with respect to their respective controls) using limma/voom, at an adjusted p-value<0.05 (see Methods). We identified a total of 339, 27, 1781 and 208 DEGs in CoV-2, RSV, H1N1 and RV16 datasets, respectively, with a more significant proportion of genes upregulated, in all cases, in response to infection (Supplementary Table 1). Our analysis indicated a significant over-representation of functional categories associated with immune response, across all infections (among upregulated genes) (Figure 1). The categories included, “response to virus”, “TNFα signaling via NFκB”, “interferon response” (types I/α, II/γ) and “apoptosis”. However, “coagulation” and “keratinocyte differentiation pathways” were among the top pathways to be uniquely enriched within the CoV-2 dataset. Genes including *MMP9, ANXA1, C1S, C3, CFB, F3, ITGB3, MAFF, MMP1, MMP9, PDGFB, PLAT, PLAU, SERPINB2, TFPI2* associated with “coagulation” and *S100A7, ANXA1, TGM1, IVL*, and small proline-rich proteins *SPRR2A, SPRR2E, SPRR2D*, associated with the “keratinocyte differentiation pathway” were among the 161 genes that were “uniquely” significantly upregulated within the CoV-2 dataset (Figure S1).

**Figure 1.**
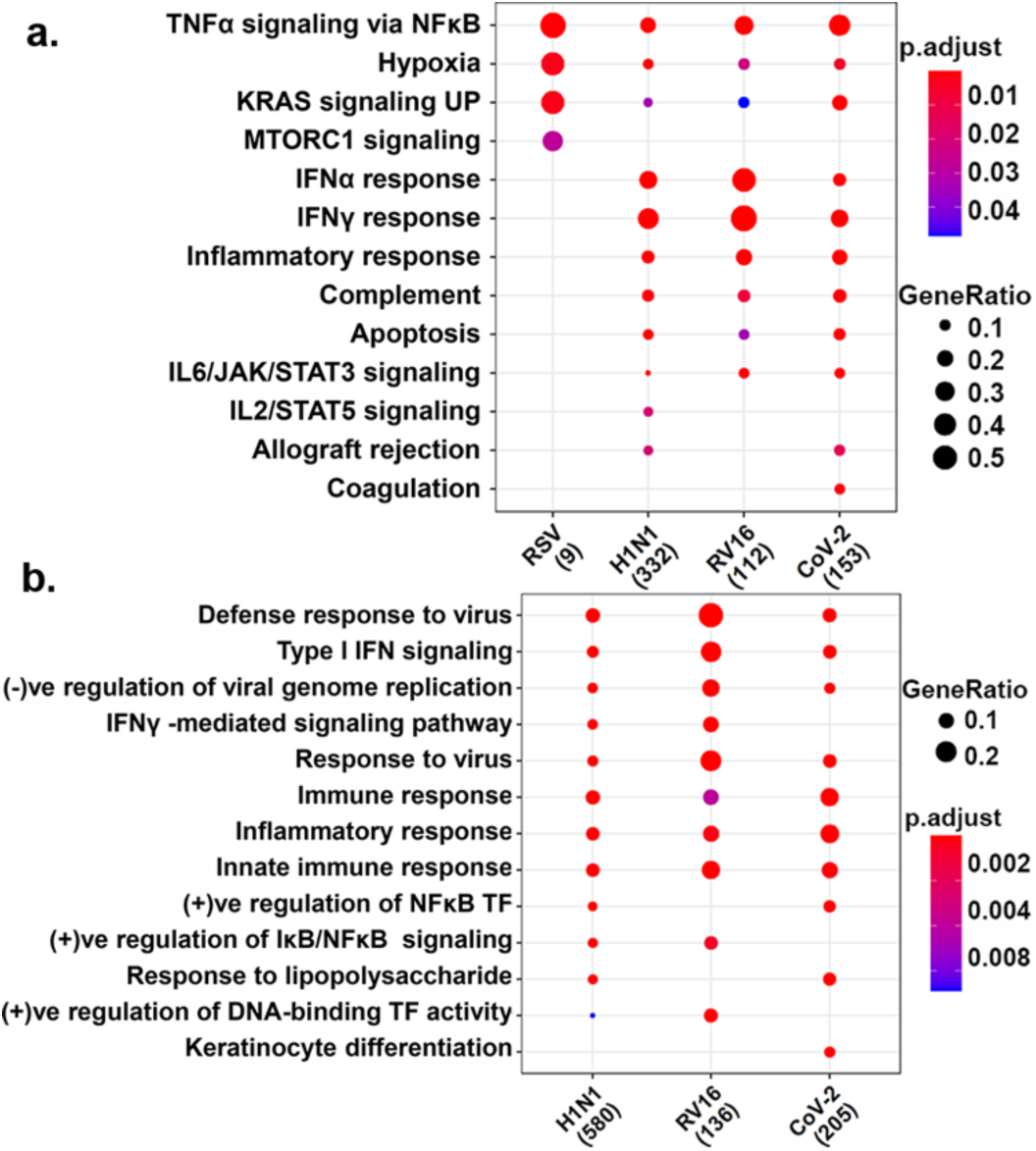
Functional enrichment of upregulated genes in upper respiratory tract infections. Functional enrichment of genes upregulated upon infection with Rhinovirus (RV16), respiratory syncytial virus (RSV), influenza (H1N1) and SARS-CoV-2 (CoV-2) were identified using **a.** mSigDB’s hallmark gene sets and **b.** Gene ontology’s (GO) biological process enrichment. No significant GO enrichment was identified for RSV and hence not included in **b.** Enrichment illustrated the biological functions common and unique to the different infections. Gene ratios are indicated by the dots’ size and the adjusted p-value is by the color scale indicated.

We posed the question if analyzing the molecular cascades associated uniquely with SARS-CoV-2, identified above, could translate to findings that are relevant to thrombotic outcomes seen in severe/acute COVID-19 patients^2^. In order to relate the clinical and molecular phenotypes associated with SARS-CoV-2 infection, we generated a protein interaction map (strength >0.85) from DEGs identified within the CoV-2 dataset (see Methods) and annotated it with data from the comparative analysis for the three other infections (RSV, RV16 and H1N1). We subsequently clustered this network, to identify “functional modules” relevant to the pathology of SARS-CoV-2 infection. Our clustering identified nine modules with five smaller functional modules containing genes associated uniquely with SARS-CoV-2, involving angiogenesis, tubulins, keratinocytes, and small proline-rich proteins (Figure 2). Three of the four remaining modules were mainly associated with immune response. These functional modules included (i) genes associated with response to IFN type I/III such as *IFITM1, IFITM3, MX1, MX2, OAS1, OAS2, IFIT1, IFIT3*, and *IRF9*; (ii) Activation of the IL-17 pro-inflammatory cascades (*IL6, IL1B*, and *CXCL1*) via the TNFα and NFκB signaling cascades (*TNF, CSF2, CSF3, NFKB1, NFKB2, RELB*, and *IRAK2*) (iii) Signaling cascades downstream of interleukin response including JAK-STAT containing genes such as *IL6R, STAT1*, SOCS3, *IFNGR1* and *PDGFB*. Interestingly, though these modules were broadly representative of vital immune mechanisms seen in all respiratory infections^21,22^, they also highlighted certain mechanisms unique to SARS-CoV-2. For example, subversion of innate immune response is inherent to cytopathic pathogens like SARS-CoV (a related beta coronavirus), which causes an increased release of pyroptosis products (e.g. IL-1B), inducing acute inflammatory responses. The antagonism of interferon response by SARS-CoV viral proteins has been suggested to occur at multiple stages of the interferon and NFKB signaling cascades, through multiple mechanisms, including IKBKE and TRAF3 regulation, affecting downstream *STAT1* associated signaling^23^. It is possible that, in the SARS-CoV-2 infection, the observed upregulation of *IKBKE, TRAF3, NFKB1, NFKB2, RELB, IRAK2* at 24hpi could be a consequence of similar mechanisms^7^ The fourth large module, contained genes specifically activated within the CoV-2 dataset, involved in fibrinolysis and plasminogen activation cascades (*PLAT, PLAU, PLAUR*, and *SERPINB2*), the complement activation cascade including *C3, CSB, CXCL5*, and platelet aggregation (platelet activation factor receptor-*PTAFR* (Figure 2)). This module was particularly noteworthy given the enrichment results (Figure 1A) and clinical findings of COVID-19^2^. It is analyzed further in the following sections.

**Figure 2.**
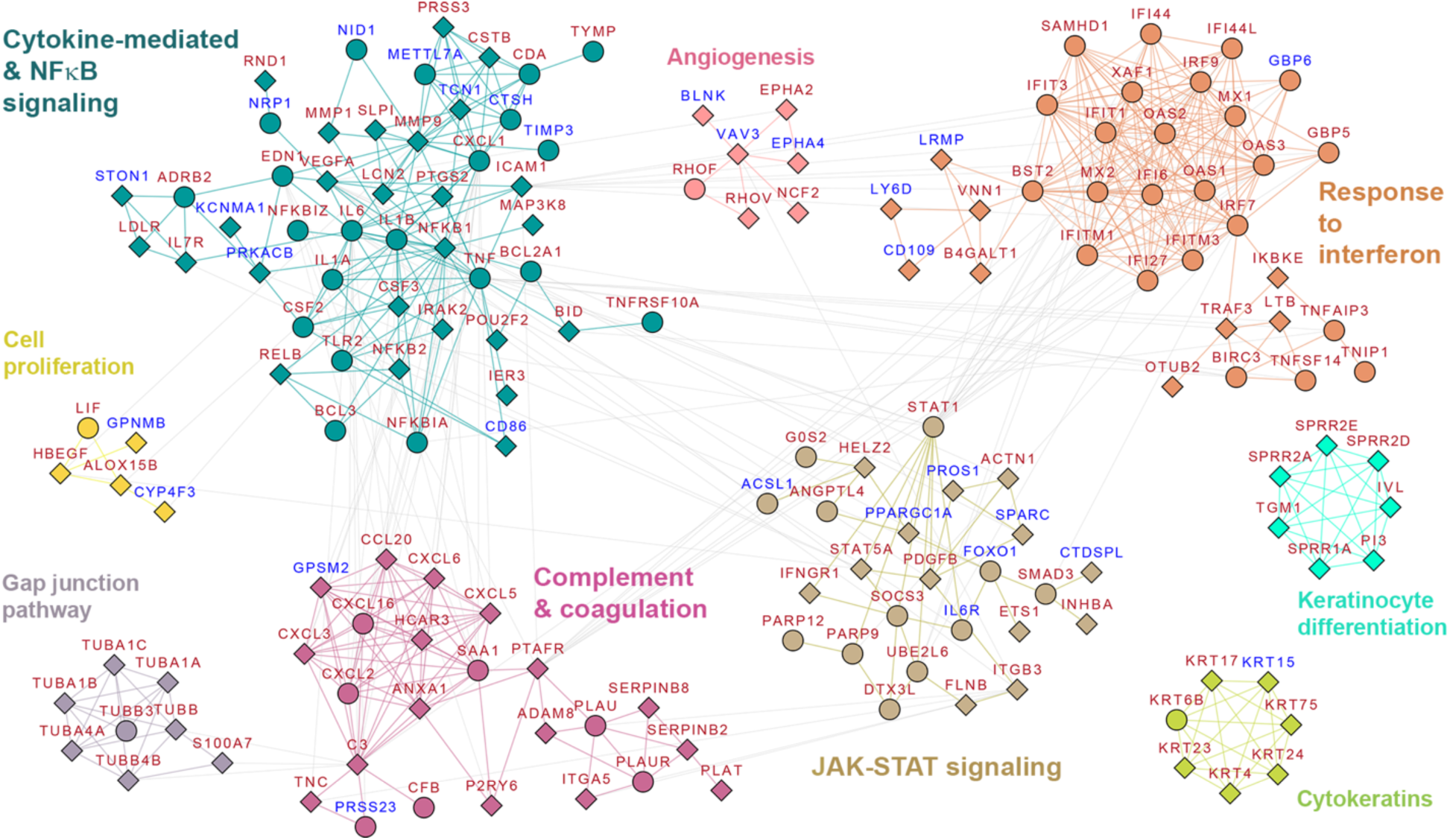
Protein interaction network for SARS-CoV-2 infection. The protein-protein interaction network extracted for genes differentially regulated (DEGs) in CoV-2 dataset (see Methods) is shown here. Clustering identified nine functional modules for further analysis. The modules functionally corresponded with i. Response to interferon, ii. Cytokine-mediated & NFkB signaling iii. Complement & coagulation iv. JAK-STAT signaling v. Cell proliferation vi. Gap junction pathway vii. Cytokeratins viii. Keratinocyte differentiation and ix. Angiogenesis. Red node labels indicate upregulated genes and blue node labels indicate downregulated genes. Diamond node shapes indicate DEGs identified only within CoV-2 while circle indicate DEGs identified in more than one upper respiratory tract infection.

Plasminogen is the precursor of the serine protease *plasmin*, a vital component of the fibrinolytic system, essential for ensuring immune cell infiltration and cytokine production^24^. Plasminogen can be activated to plasmin by two serine peptidases, the tissue tissue-specific (T-PA) and urokinase (U-PA) plasminogen activators encoded by *PLAT* and *PLAU*, respectively. It has been previously reported for SARS-CoV and influenza infections that dysregulation of the urokinase pathway including U-PA(*PLAU*) and its inhibitor PAI-1*(SERPINE1)* might contribute to the severity of lung disease by altering the dynamics of fibrin breakdown and intra-alveolar fibrin levels and subsequently inflammation^25,26^. Here we observed a similar dynamic with SARS-CoV-2 infection albeit through activation of tissue plasminogen, *PLAT*, and the inhibitor *SERPINB2*. T-PA (which is triggered when bound to fibrin) and *F3* (tissue factor, activated in CoV-2) levels are known to correlate with d-dimer levels^27^. D-dimer, a product of fibrin degradation by plasmin, is elevated in patients with COVID-19 and has been identified as a marker for disseminated intravascular coagulopathy and a worse patient prognosis^28^. These findings indicate increased fibrinolytic activity, specifically via T-PA (*PLAT)* activation, in SARS-CoV-2 infections.

The outcome of any viral infection is mediated through a complex interplay between viral and host proteins, which allow for a coordinated innate immune response. Plasminogen inhibitor (PAI)-1 (*SERPINE1*) has been reported to function as an anti-viral factor capable of inhibiting extracellular maturation of influenza particles, specifically through their action on *TMPRSS2*^29^. A similar mechanism involving PAI-2 (*SERPINB2)* likely exists in SARS-CoV-2 infections. Additionally, T-PA (*PLAT*) has been reported to interact with *ORF8* protein of SARS-CoV-2 virus^7^; however, its consequence has not yet been elucidated. We hypothesize that if this interaction titrates out T-PA, it is likely to result in reduced levels of plasmin which could contribute to downstream thrombosis events, as seen in COVID-19 patients^2^. Increased expression of *PLAT* validated this. Additionally, increased concentration of *SERPINB2* is also likely to reduce plasmin levels leading to thrombosis. The unique regulation of *SERPINB2* and *PLAT* in SARS-CoV-2 infections and their subsequent effect on viral-host interaction dynamics is worthy of further investigation.

Matrix metalloproteinases (MMPs) are a family of proteolytic enzymes that facilitate leukocyte infiltration by breaking down the extracellular matrix (ECM) and basement membrane. The genes *MMP1* and *MMP9* (both upregulated specifically within the CoV-2 dataset, see Figure 2) can also activated by plasmin *in vivo* and *in vitro*^30^. Additionally, degranulation of neutrophils by T-PA has been indicated as a source for increase of matrix metalloproteinase 9 (MMP-9)^31^. Given the observed increase in MMPs, and their role in facilitating lung inflammation in ARDS (by enabling neutrophil migration and ECM breakdown)^32^, it will be essential to evaluate the impact of T-PA treatments^33^ on pulmonary remodeling (via MMPs) in patients with severe/acute COVID-19.

We identified significant upregulation of several genes encoding components of the complement system, including *C3, CFB* and *C1S*, uniquely within the CoV-2 dataset, in contrast to other upper respiratory tract infections (at 24 hpi). The complement pathway is an integral part of the innate immune response and is involved in immunosurveillance for pathogen clearance (bacterial and many viral)^34^. It traditionally serves as a vital link between the innate and the adaptive immunity, driving pro-inflammatory cascades. Several of the chemokines/chemoattractants (*CXCL5, CXCL6, CXCL3*, and *CCL20*) identified within this cluster can be also be activated by signaling events precipitated by complement activation^35^. Overstimulation of these chemokines, particularly CXCL5, is known to cause destructive inflammatory lung condition in certain pathogenic models of lung disease^36^.

Plasmin also activates the complement cascade independently of established pathways (alternate, lectin, and classical pathways) by cleaving *C3* and *C5* to functional C3a and C5a respectively, both of which are known to be crucially involved in the inflammatory response^37^. Furthermore, there is increasing evidence for the role of complement in coagulation. C3 is shown to bind fibrinogen and fibrin with high affinity and prolonging fibrinolysis in a concentration dependent manner^38^. Additionally, studies in animal models of thrombosis have identified a plasmin driven C5a generation capable of driving procoagulant cascades^39^. Moreover, the presence of terminal complement components includes C5b-9 (membrane attack complex), C4d, and mannose-binding lectin (MBL)-associated serine protease (*MASP2*) within the pulmonary microvasculature and purpuric skin lesions of deceased COVID-19 patients^5^. These findings further provide evidence for the activation of complement components observed here and their role in plasmin-mediated thrombotic events, as a likely mechanism in COVID-19.

Unique activation of the platelet-activating factor (PAF) receptor (PAFR, encoded by the gene *PTAFR*) within the complement and coagulation cluster was particularly interesting (Figure 2). PAF is a potent activator of platelet aggregation as well as immune cell types including macrophages and neutrophils^40^. PAF binding to PAFR activates several intracellular signaling events, and plays pathophysiological roles in molecular mechanisms underlying anaphylaxis, bronchial asthma, cystic fibrosis^41^, and endotoxin shock/sepsis^42^. It is a crucial mediator of systemic and pulmonary hemodynamic changes. Immune infiltration primed by interaction between complement system and PAF has been reported previously. Specifically, C5b can interact with PAF to bind PAFR on the lung epithelial cells increasing vascular permeability^43^, and inducing massive eosinophil transmigration^44^. Eosinophils release leukotrienes and lipids which can cause further epithelial damage and airflow obstruction^45^. Leukotrienes, C5a and PAF are potent chemoattractants of polymorphonuclear leukocytes (neutrophils or PMNs) which can further activate neutrophil extracellular traps (NETs) leading to pulmonary damage seen in COVID-19 patients^46^. Given the ubiquitousness of SARS-CoV-2 receptors and its ability to infect a broad range of cell types^18^, analysis of *PTAFR* activation patterns in infected tissues (including vascular endothelia cells) could provide further insights into increased platelet aggregation (via PAF signaling) and the risk for arterial thrombosis seen in severe/acute COVID-19 patients ^2^. Vitamin-D is known to attenuate *ICAM1* (a potent activator of inflammation and epithelial-mesenchymal transition) and *PTAFR* expression and subsequently NFκB mediated inflammation, in rhinovirus infections^47^. Similar mechanisms can possibly underly the recent reports on reduced risk for COVID-19 infections after Vitamin-D supplementation, and provide evidence for PAF/PAFR mediated signaling in the progression of COVID-19^48^.

The significant enrichment of mechanisms identified above were indicative of active epithelial remodeling and provided evidence for a plasmin-mediated complement activation within SARS-CoV-2 in contrast to other respiratory infections. We also identified mechanisms that might contribute to increased inflammation via immune infiltration and platelet aggregation, primed by interactions between complement system, plasmin and the PAF/PAFR. Our findings further the understanding of modulation of hemostatic factors, affecting the mechanics of circulating fibrin and fibrin breakdown products^8^ and complement activation, underlying acute thrombotic events including arterial thrombosis in COVID-19 patients^2^. Although we cannot wholly ascribe all mechanisms observed within an *in vitro* study as contributing to observed clinical pathology of COVID-19, our analysis highlighted SARS-CoV-2 driven response of healthy lung epithelial cells and its ability to incite a misdirected cascade of interactions between the coagulation and complement activation, mediated by plasmin. In addition, analysis of DEGs from transcriptomic measurements in postmortem lung tissue from COVID-19 patients provided a partial insight into the mechanisms associated with virus-altered epithelial function. Functional analysis of DEGs identified with COVID-19 exhibited an upregulation of similar immune cascades as those identified within the CoV-2 dataset (Figure S2 and S3). Significant correlation was also identified for foldchanges between DEGs commonly regulated between COVID-19 lung and CoV-2 dataset (rho=0.34). We present our current view of the crosstalk between plasminogen, complement and platelet-activating systems in SARS-CoV-2 infection in Figure 3.

**Figure 3.**
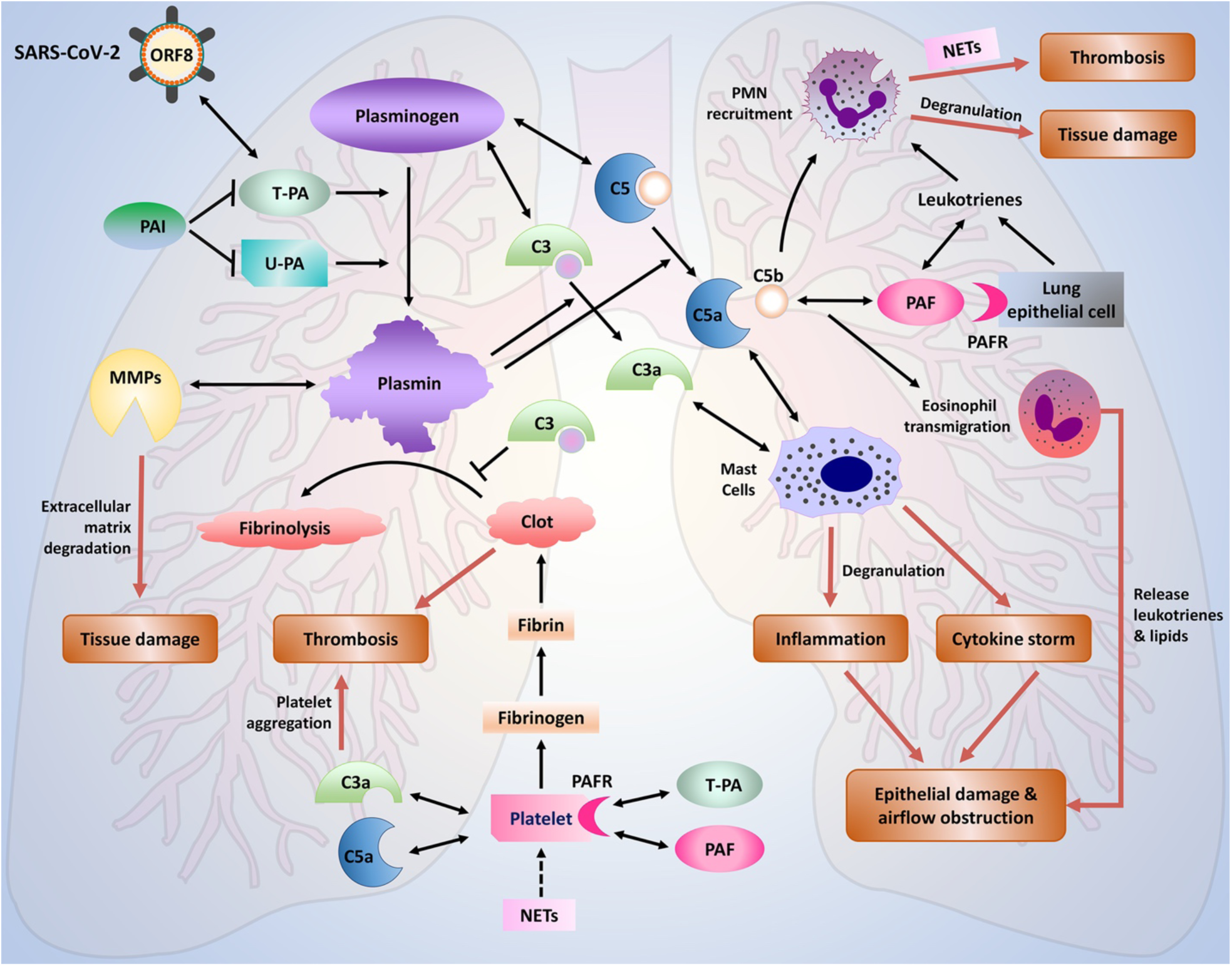
A schematic representation of the crosstalk between plasmin, complement, and platelet-activating systems in SARS-CoV-2 infection. Plasminogen conversion is mediated by either by tissue-specific plasminogen activator (T-PA) or urokinase plasminogen activator (U-PA), whose activities can be inhibited by the inhibitors, PAI-1 (SERPINE1) or PAI-2 (SERPINE2B). The conversion of plasminogen to active plasmin is critical for blood clot breakdown. Failure to breakdown the clots (fibrinolysis) leads to thrombosis. Fibrinolysis can be inhibited by complement component C3 (C3). The complement components C3 and C5 can be activated by plasmin (extrinsic pathway), in addition to intrinsic (classic, lectin, and alternative) pathways. The anaphylatoxins, C3a and C5a, interact and stimulate mast cells to degranulate, releasing histamine, cytokines, granulocyte-macrophage colony-stimulating factor, leukotrienes, heparin, and several proteases that damage the tissues. Overstimulation of complement cascade leads to inflammation, cytokine storm resulting in epithelial damage, and airflow obstruction that manifests as the acute respiratory distress syndrome (ARDS). Also, C5b and Leukotriene-bound PAF are potent attractants of polymorphonuclear leukocytes (PMNs) to the site of damage. The recruited PMNs can release microbiocidal molecules and form the neutrophil extracellular traps (NETs). NETs are proinflammatory and promote tissue damage, thrombus formation, and activate platelets. Degradation of the basement membrane/ ECM promote by matrix metalloproteinases further promote immune cell infiltration and tissue damage. NETs, tissue injury, platelets-activating factors (PAF), T-PA (if overexpressed), C3a and C5a activate platelets to aggregate on a fibrin scaffold to form clot. Clots and tissue injury due to these mechanisms lead to airflow obstruction that manifests as ARDS. Double arrows indicate known interactions.

Further *in vivo* studies will be required to better understand the viral-host dynamics with respect to T-PA (*PLAT)* and PAFR (*PTAFR)* activation and the subsequent systemic inflammatory responses, both in symptomatic and asymptomatic populations. We believe that managing the equilibrium between the complement, coagulation and platelet-activating components can determine the overall biological activity and the outcome of a disease severity in COVID-19. Investigation into the efficacy of complement inhibitors (such as Eculizumab^49^ or Compstatin^50^), and plasminogen activators specifically for T-PA, combined with Vitamin-D supplementation to alleviate symptoms associated with systemic thrombosis and ARDS, should be pursued in patients with severe/acute COVID-19.

## ACKNOWLEDGMENTS

Supported by the National Institute of Health Grants U19 AI090023(SS) and R01 LM012595(SS).

## AUTHOR CONTRIBUTIONS

KM and SS conceived the study and designed the work, KM carried out data acquisition and the analyses, and interpreted the data with help from KMa and SS, SS supervised the study, and KM developed the first draft of the manuscript which was revised by all authors.

## ONLINE METHODS

### Data acquisition

We analyzed the data recently published by Blanco-Melo et al^51^ to gain mechanistic insights into the pathogenicity of SARS-CoV-2. The published dataset contained multiple cell lines treated with SARS-CoV-2 including NHBE, Calu-3, A529 (with and without exogenous expression of ACE2) in addition to COVID-19 lung and normal tissue samples (available via GSE147507). Histogram of counts within normal and diseased lung samples indicated that one COVID-19 lung sample (Covid Lung 2, Figure S4) is a likely an outlier and was ignored from further analysis. Based on the sample clustering results of the raw counts (Figure S2.B), we limited our analysis to NHBE/ normal human bronchial epithelial cell lines (hence forth referred to as the CoV-2 dataset). All available series (GSE) in GEO were extracted from the gene expression omnibus with key words-“SARS-CoV”, “MERS-CoV”, “RSV” or “respiratory syncytial virus”, “Influenza” and “Rhinovirus” for respiratory cell lines (NHBE or BEAS-2B). Since the CoV-2 transcriptional data was processed at 24 hpi (hours post infection), we chose to compare only those infections which had cell-lines at 24 hpi yielding the following series GSE3397, GSE71766, GSE100504, GSE81909, GSE27973, and GSE28904. To limit the impact of sequencing technologies, we utilized only Affymetrix, the technology with the most coverage among the series considered. Applying the above stringent inclusion criteria yielded 3 GEO series GSE3397^52^ (RSV), GSE71766^53^ (Influenza/H1N1 and Rhinovirus/RV16) and GSE27973^54^ (Rhinovirus/RV16). We, however, did not find studies on related beta-coronaviruses in NHBE/BEAS-2B cell lines which matched our inclusion criterion.

### Differential expression analysis

For the sake of reproducibility, we called differentially expressed genes (DEGs) at adjusted p.value <0.05, using GEO2R for GSE3397, GSE71766 and GSE27973 comparing infected cells with their respective mocks at 24 hpi only. The DEGs called for GSE27973 were a subset of GSE71766 RV16 comparisons and subsequently ignored. Since these were microarray studies, we aggregated probes which were significantly differentially expressed, to gene names and calculated a mean fold change (Supplementary Table 1) and considered these for all comparisons outlined in the manuscript. We utilized the same pipeline implemented in GEO2R to call DEGs from CoV-2 data (using the limma-voom pipeline in R). Low counts were filtered using the “FilterByExpression” feature available through the edgeR package. “topTable” was used to extract all samples under our significance threshold of p.adj <0.05.We would additionally like to point out that we reanalyzed the CoV-2 data using the DESeq2 protocol as described in the original publication consistently identified similar number of DEGs as detected through limma-voom. For the comparision of COVID-19 lung biopsy sample against two healthy tissue biopsies, we utilized the exact T-test via “edgeR” to establish significance and extracted the significant genes via “topTags”.

### Protein network construction

The human protein-protein interaction network (PPIN) was downloaded from STRINGdb (v 11.0)^55^ for a combined edge strength of >0.85. We extracted a SARS-CoV-2 infection relevant subnetwork from this PPIN using DEGs identified in the CoV-2 dataset. The resulting CoV-2 network contained 272 nodes and 608 edges (Figure S3). We additionally annotated this CoV-2 network with differential gene expression information (foldchange) identified in all infections, if present, to allow us to identify DEGs unique to CoV-2. Functionally relevant modules were extracted using the clusterMaker plugin in Cytoscape^56^. Clusters with >3 nodes were retained for further analysis, resulting in a network size of 162 nodes and 9 clusters (Figure 2). The clusters were named based on the most prominent terms associated with each cluster as identified using Enrichr^57^. Functional enrichment was performed using gene ontology (biological process) and mSigDB’s Hallmark genesets (v7.1). All visualizations were generated via the ClusterProfiler library^58^ available through R/Bioconductor.

## SUPPLEMENTARY FIGURES

**Figure 1.**
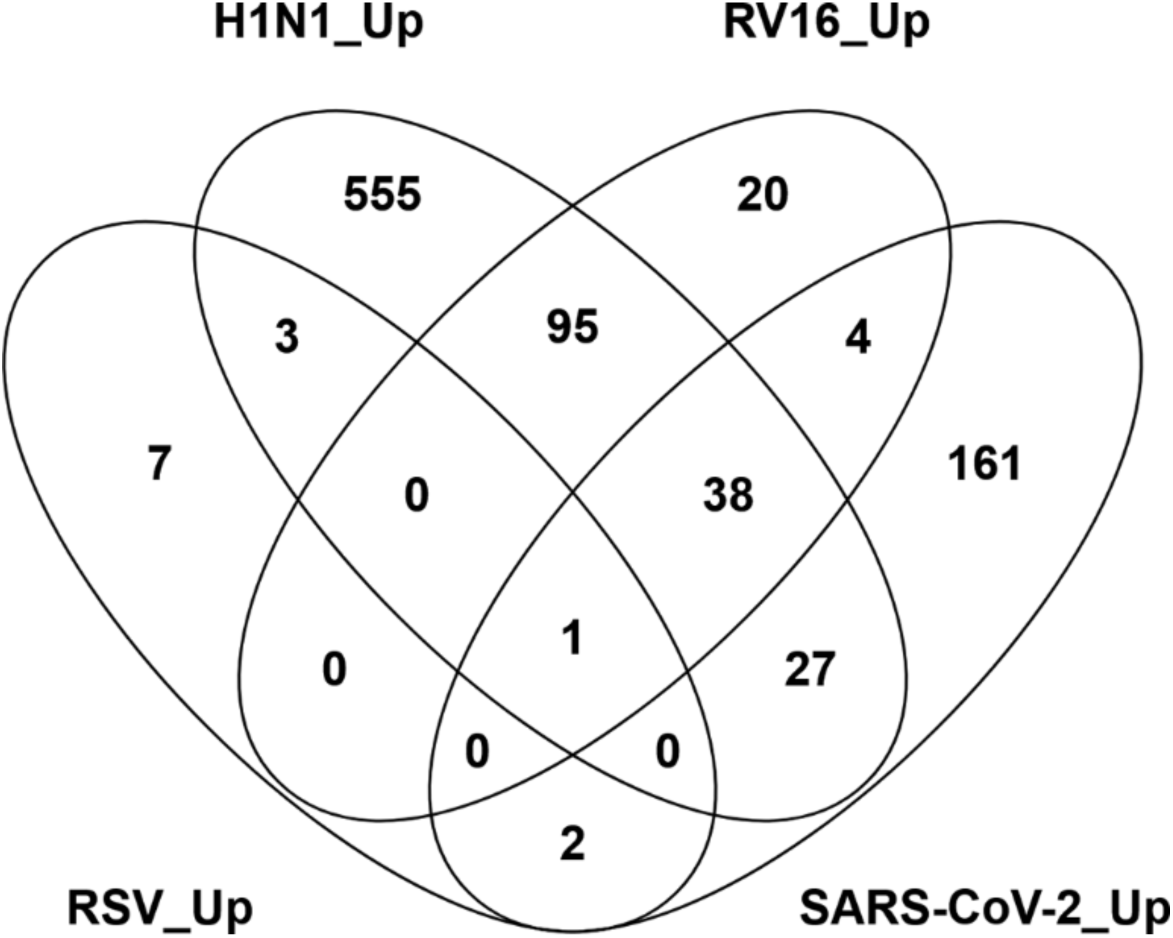
Overlap of upregulated gene across four upper respiratory tract infections (Rhinovirus-RV16, respiratory synytical virus RSV, and influenza H1N1) and SARS-CoV-2 (CoV-2), compared to their respective control, represented as a Venn diagram. 161 genes are uniquely differentially upregulated in SARS-CoV-2 infected cells.

**Figure 2.**
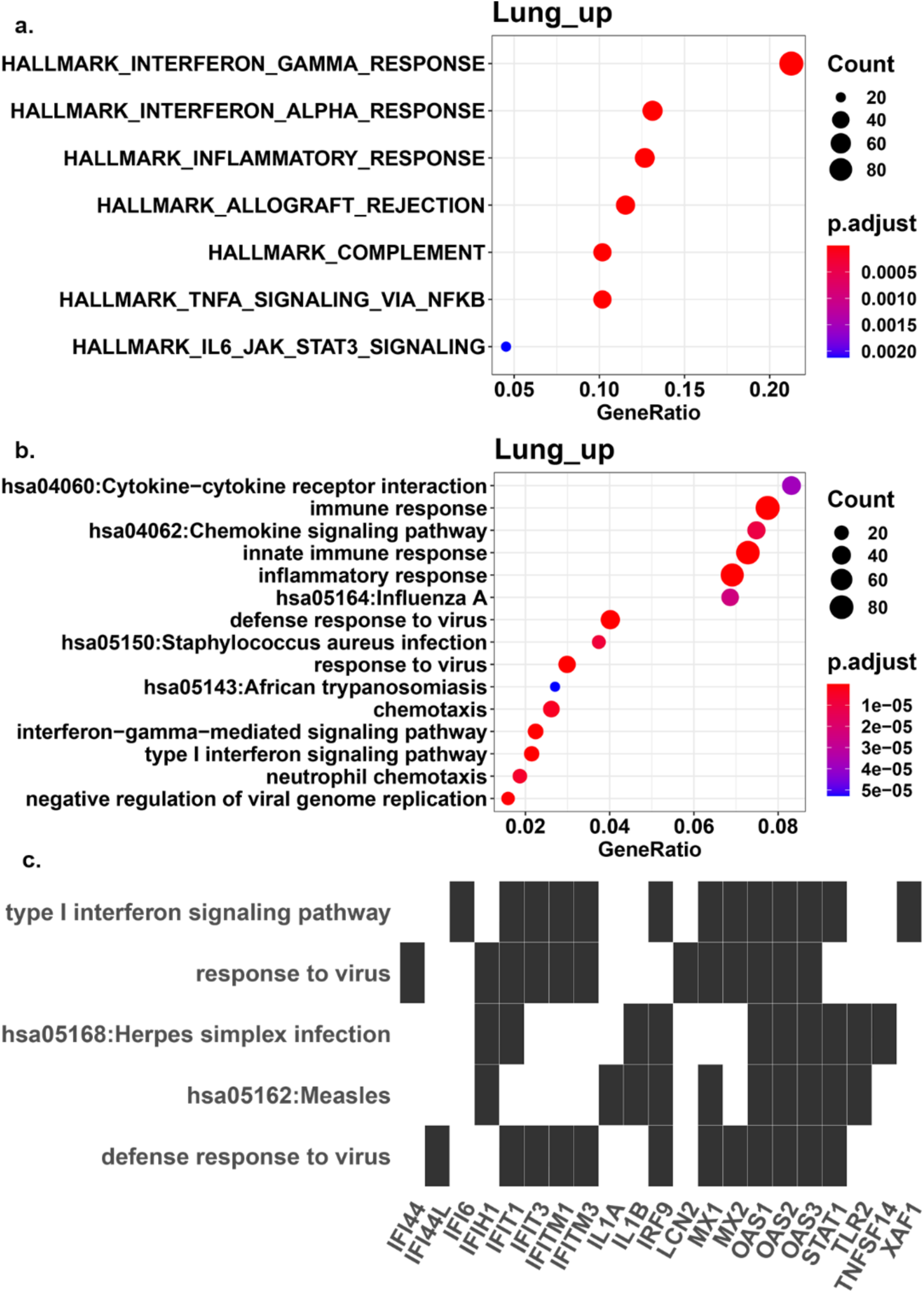
Functional analysis of differentially expressed genes (DEGs) identified in COVID lung. Functional enrichment was identified for genes upregulated in COVID-19 lung tissue using a. mSigDB’s hallmark gene sets and B. Gene ontology’s biological process enrichment c. A heatmap plot showing the functional enrichment of genes which overlap between DEGs upregulated in CoV-2 dataset and COVID-19 lung. Number of genes within the dataset for each category are represented by the size of the dots and the p-adjusted value by the color scale indicated.

**Figure 3.**
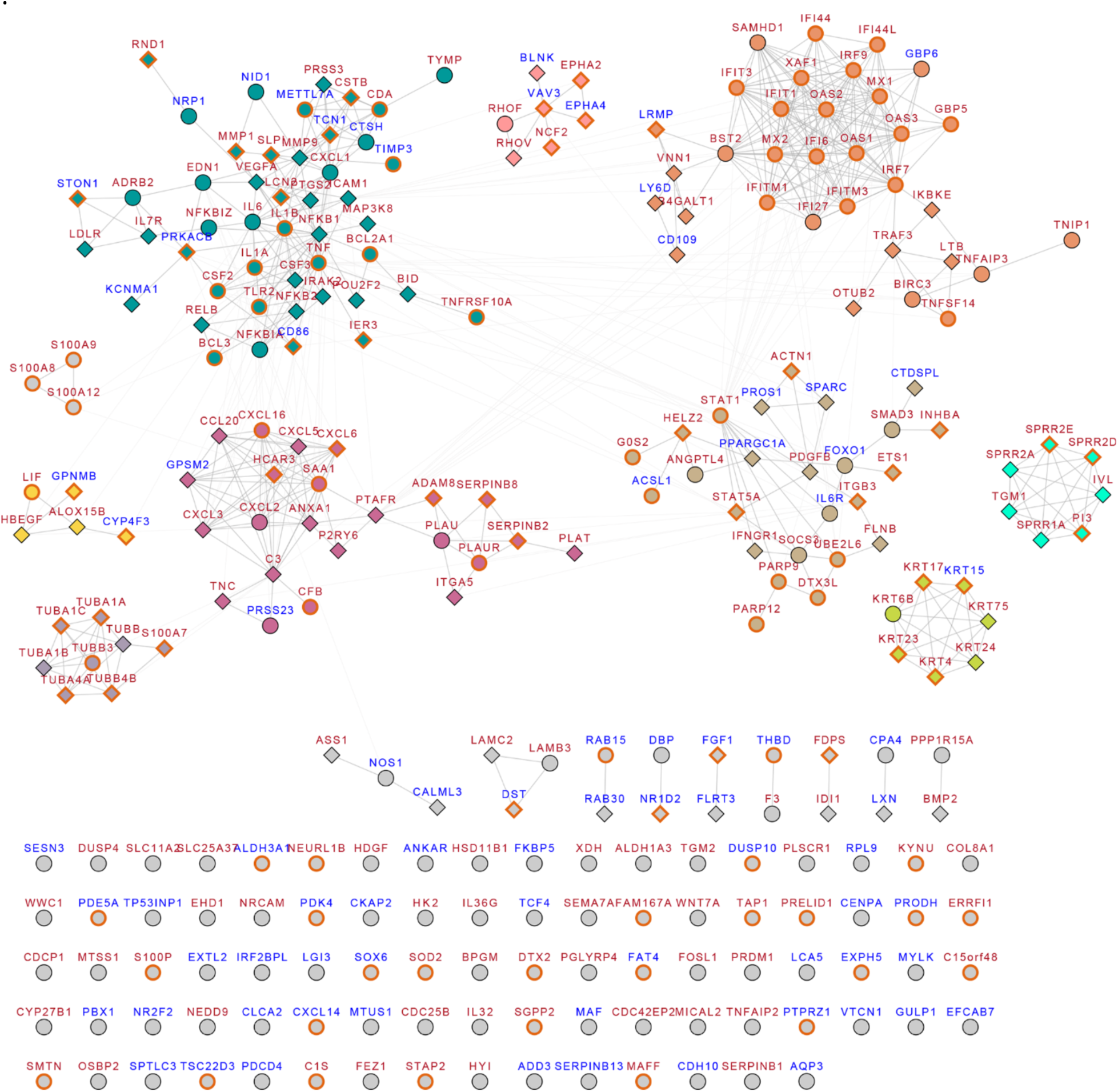
Protein interaction network for SARS-CoV-2 infection-The entire protein-protein interaction network extracted from STRINGdb for genes differentially regulated (DEGs) in CoV-2 dataset (see Methods) is shown here. All nodes with n>3 were extracted and functionally annotated and representd in Figure 2. This figure additionally highlights genes that were identified as DEGs within the COVID-19 lung samples (indicated with an orange node border). Red node labels indicate upregulated genes and blue node labels indicate downregulated genes. Diamond node shapes indicate DEGs identified only within CoV-2 while circle indicate DEGs identified in more than one upper respiratory tract infection.

**Figure 4.**
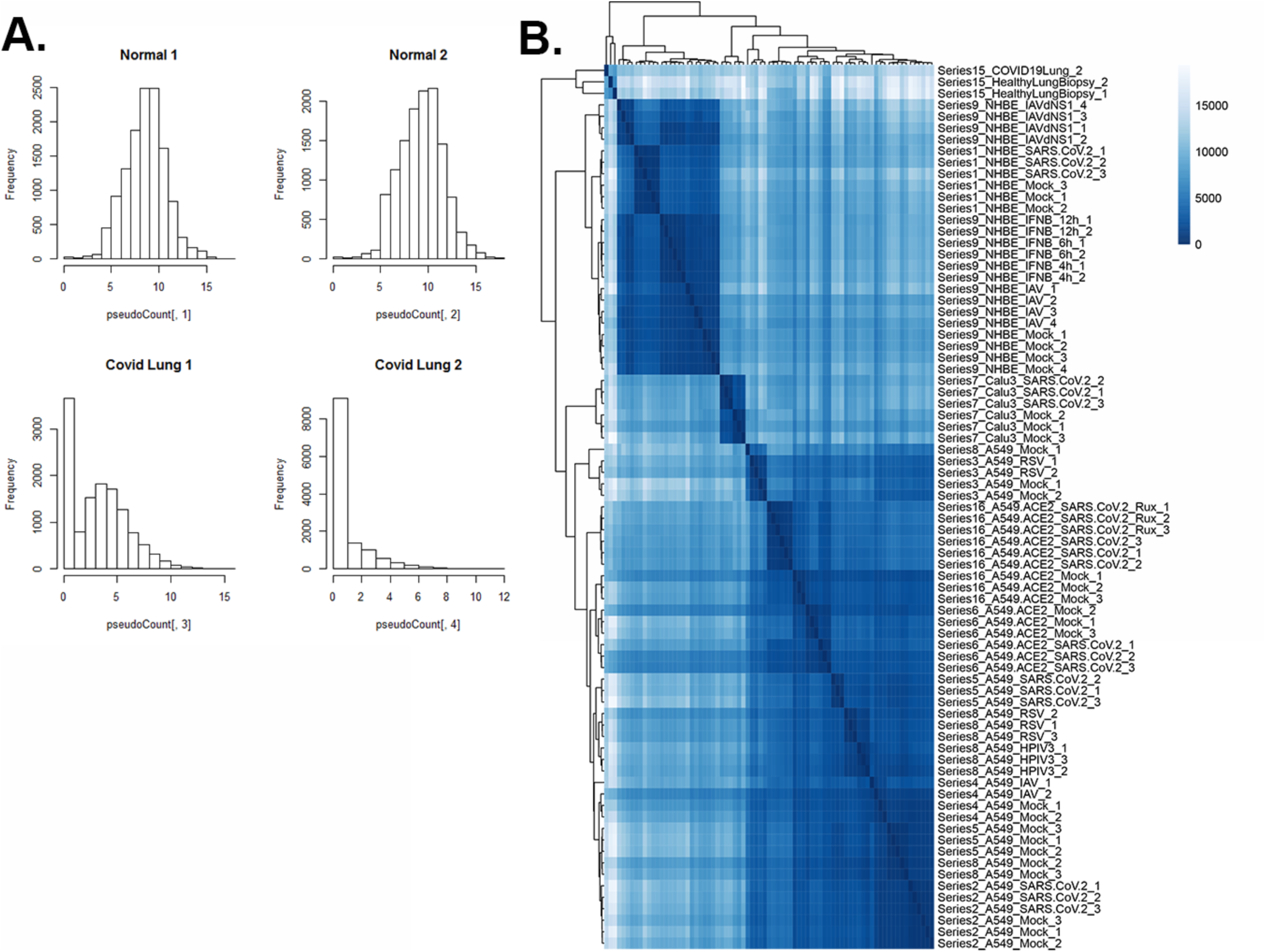
Analysis of previously published SARS-CoV-2 sequencing data and rationale for using NHBE cell lines and only one COVID-19 lung tissue A. Histograms of raw counts (log 2 scale, x axis) as published in GSE145708 for biopsies obtained from normal and COVID-19 lung (n=2). These results indicate likely degradation of covid lung 2 samples. We subsequently consider this sample unusable for downstream analysis of fold changes between healthy and COVID lung B. Clustering of counts across all cell lines infected with SARS-CoV-2, Influenza (and their respective controls) and one lung biopsy as obtained from the original study. Based on the clustering tree, we reasoned that the NHBE cell line better approximates lung biopsies and used them as the primary cell line of interest for this analysis

